# Differential sex-dependent responses of circulating steroid hormones and cortical gene expression in a preclinical traumatic brain injury model

**DOI:** 10.64898/2026.02.06.703864

**Authors:** Amberlyn Simmons, Sierra F. Wilferd, Sofia Campagnuolo, Veronica Pena, Heather Bimonte-Nelson, Jason Newbern, Rachael W. Sirianni, Christopher L. Plaisier, Sarah E. Stabenfeldt

## Abstract

Accumulating evidence supports sex differences in traumatic brain injury (TBI) outcomes, however the underlying processes that lead to sex differences are not well understood. TBI results in the initiation of molecular and cellular responses that facilitate the progression of neurodegeneration. Importantly, little is known about how the circulating hormone profile is altered in response to TBI, and whether sex differences in endocrine responses might shape secondary injury pathologies. Using intact male and female mice in a preclinical TBI model, we assessed changes in plasma hormone concentrations and cortical gene expression at 24 and 72 hours after TBI. We demonstrate that males and females exhibit sex-specific alterations in circulating levels of progesterone, testosterone, androstenedione, estradiol and dehydroepiandrosterone (DHEA) in response to TBI. We also identified sex differences in the expression of genes that are involved in immune responses and tissue remodeling after injury. Moreover, we report divergent circulating hormone and gene expression correlations between sexes.

## Introduction

Traumatic brain injury (TBI) is a major cause of death and disability worldwide, with approximately 70 million individuals sustaining a TBI globally each year (1). Within this vast patient population, the incident rates for TBI differ between male and female individuals, where males have higher rates of TBI-related hospitalizations and deaths compared to females (2).

Beyond TBI incidence rates, clinical outcomes and persistence of symptoms post-TBI differ between males and females. Clinical studies reveal females report worse outcomes including significantly higher post-concussion symptom scores and worse six-month outcomes compared to males after injury (3–5). Given these clinical findings, it is critically important to examine sex as a biological factor in preclinical models and clinical assessments to gain a deeper understanding of mechanistic drivers that contribute to differential responses in TBI outcomes.

TBI induces dynamic changes in circulating steroid hormones that may modulate the evolution of secondary injury pathologies (6). Injury-induced stress signals cause robust activation of the hypothalamic-pituitary-adrenal (HPA) axis, leading to hormonal dysregulation (7,8). Previous clinical and preclinical studies have reported links between alterations in hormone levels and blood-brain barrier (BBB) dysfunction, neuroinflammation, and excitotoxicity (9–11). Importantly, despite known baseline differences in hormone profile between sexes, little is known about whether TBI alters circulating hormone profile differentially based on sex, nor how these potential sex-dependent changes may impact secondary injury mechanisms.

Here, we assessed murine sex-associated differences in circulating steroid hormones and cortical gene expression in response to TBI. We then conducted a subsequent correlation analysis between hormone level changes and biological processes involved in secondary brain injury. Finally, to link the transcriptomic findings to protein expression, we probed the expression of crucial BBB tight junction and adherens junction proteins within the injured cortex acutely after TBI. Together, our data provide evidence of distinct sex-specific correlations between circulating injury-induced sex steroid hormone level changes and secondary TBI pathogenesis.

## Results

### Evaluating sex-based differences in the response to TBI

We and others have observed sex differences in the response to TBI (12–15). However, the molecular mechanisms underlying differential sex-dependent alterations are not well understood. To directly evaluate sex-based responses to TBI, our experimental cohorts consisted of naïve intact male and female mice to capture baseline and male and female mice at 24 or 72 hours (hrs) post-TBI (unilateral controlled cortical impact model, CCI; Figure 1A). At each timepoint, mice were sacrificed, and blood plasma was analyzed via mass spectrometry to quantify steroid hormone levels for five analytes (progesterone, testosterone, androstenedione, estradiol, and DHEA; Figure 1B). Transcriptomic analysis (RNA-seq) on cortical punches from the injured ipsilateral and the uninjured contralateral sides of the cortex (Figure 1C) was conducted on statistically representative groups for each cohort (Figure 1A). We then performed a correlation analysis between the transcriptomic and hormone profiling data sets to gain insight into potential drivers of sex-based TBI pathology. Finally, we used immunohistochemistry to assess the protein expression of key BBB proteins that were significantly upregulated in transcriptomic analysis (Figure 1D). Collectively, the experimental design captured snapshots of multiple physiological systems at different points after TBI to assess the holistic response to TBI.

**Figure 1.**
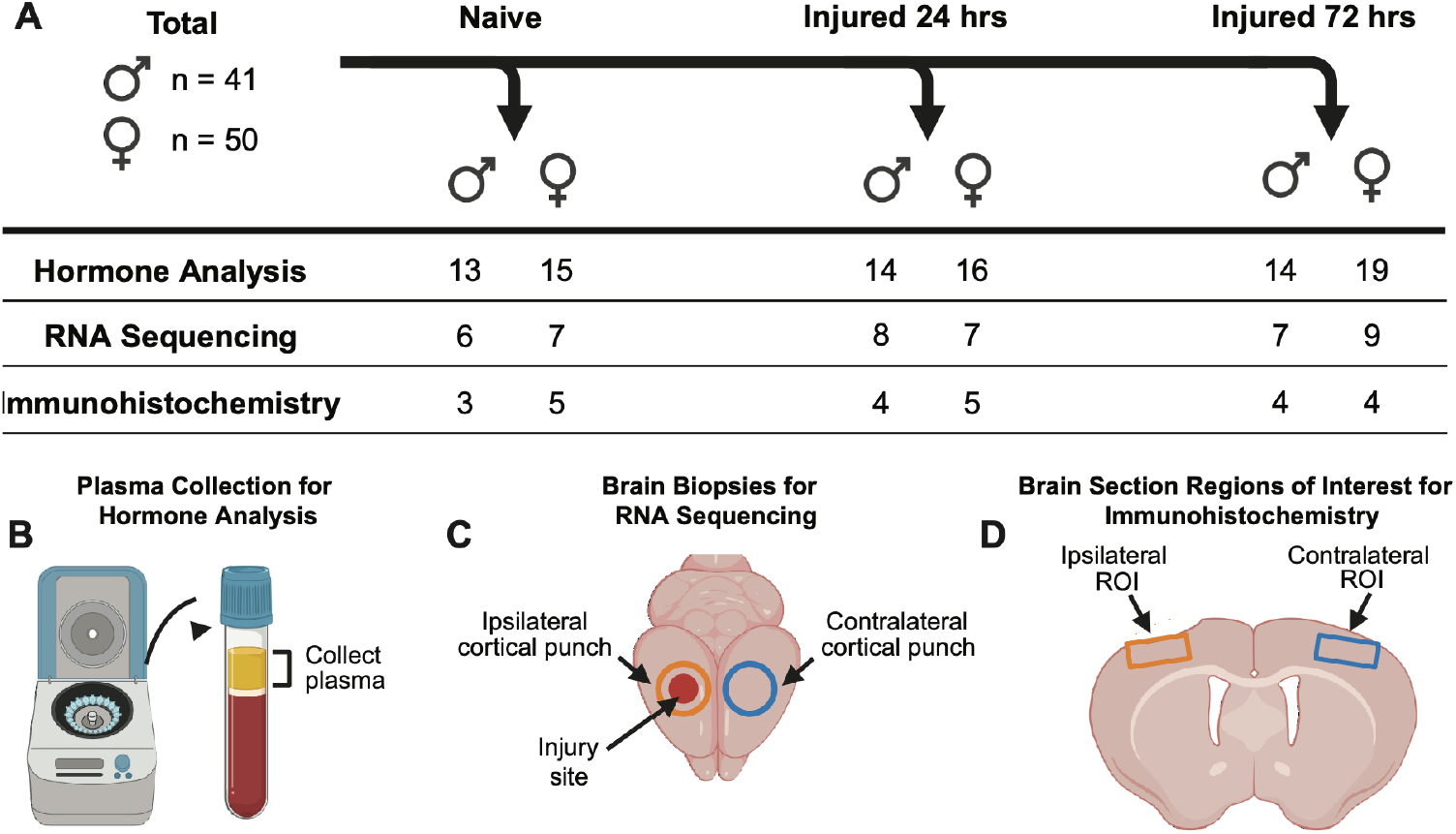
Study design and methods. Experimental design for the study including cohort description, sample sizes and analysis methods performed (**A**). Plasma was collected for steroid hormone quantification using mass spectrometry (**B**). Cortical biopsy punches were processed for RNA-sequencing analysis (**C**). Whole brains were processed for immunohistochemical analysis of ipsilateral and contralateral blood-brain barrier junctional protein expression (**D**).

### Sex-specific alterations in circulating sex steroid hormone profile after TBI

At the outset, we acknowledge that circulating sex hormone levels in adult intact females fluctuate with respect to the estrous cycle phases, and may impact neutrotrauma, pathology, behavioral outcomes? (be specific) (citation). We conducted estrous cycle tracking on our female cohorts (Supplemental Figure 1). However, the difficulty and invasiveness of repeated cycle staging, the brief duration of certain phases, and the large number of animals that would be required to adequately power comparisons across all estrous stages made it infeasible to design the experiment around cycle phase as a biological variable for our TBI studies. Instead, we randomized timing of TBI with respect to the estrous cycle phase, mimicking clincal incidence of TBI.

Mass spectrometry analysis of five sex steroids in blood plasma was completed for naïve, 24 hrs, and 72 hrs post-TBI cohorts. Significant differences among specific sex steroid hormones after TBI were determined via one-way ANOVAs across timepoints within each sex (Supplemental Table 1,2). Female mice exhibited a main effect of injury for progesterone (F=4.714, df =49, p=0.0136), testosterone (F=6.858, df=49, p=0.0024), and androstenedione (F=8.495, df=49, p=0.0007) (Fig. 2 A-C). At 24 hrs post-TBI in the female cohort, the following hormones decreased compared to naïve females: progesterone (t=2.664, df=47, p=0.0316), testosterone (t=3.139, df=47, p=0.0088), and androstenedione (t=4.010, df=47, p=0.0006). Similarly, at 72 hrs post-TBI in females, we observed decreased progesterone (t=2.719, df=47, p=0.0274), testosterone (t=3.343, df=47, p=0.0049), and androstenedione (t=2.957, df=47, p=0.0145). Conversely, male mice exhibited a main injury effect on DHEA (F=4.636, df=40, p=0.0158) and estradiol (F=4.934, df=40, p=0.0124) (Fig. 2 I,J). Males did not exhibit significant alterations in hormone profile at 24 hrs post-TBI for any measured hormone. At 72 hrs post-TBI males, we observed decreased estradiol (t=3.125, df=38, p=0.0102) and increased DHEA (t=2.941, df=38, p=0.0166) compared to naïve cohort.

**Table 1.**
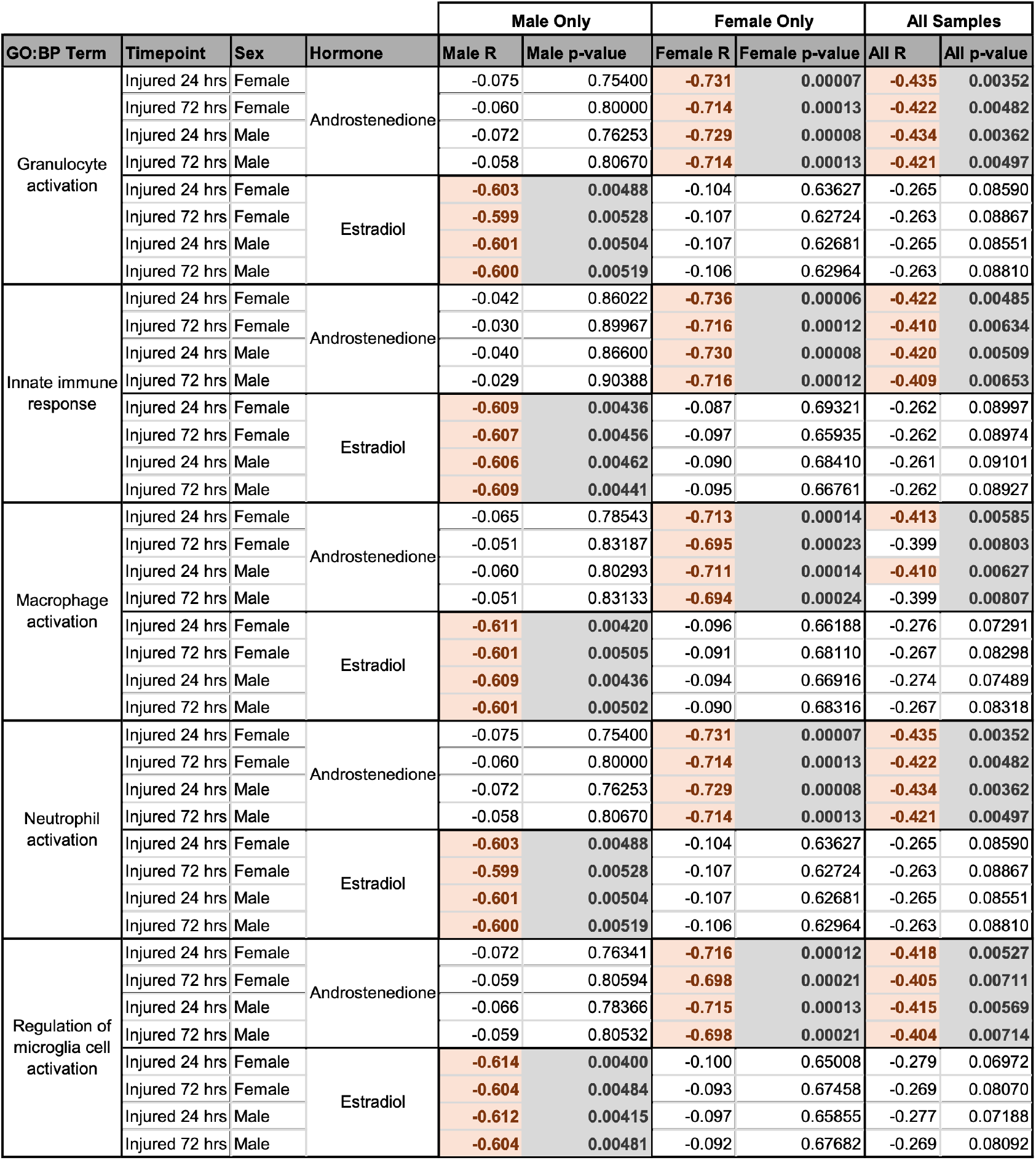
Results of Pearson correlation between eigengene vectors of genes that showed significant semantic similarity to innate immune GO:BP terms and hormone levels for each cohort.

**Table 2.** Results of Pearson correlation between eigengene vectors of genes that showed significant semantic similarity to tissue remodeling GO:BP terms and hormone levels for each cohort.

| GO:BP term | Timepoint | Sex | Hormone | Male Only |  | Female Only |  | All Samples |  |
| --- | --- | --- | --- | --- | --- | --- | --- | --- | --- |
|  |  |  |  | Male R | Male p-value | Female R | Female p-value | All R | All p-value |
| Adherens junction organization | Injured 72 hrs | Female | Androstenedione | -0.016 | 0.94522 | **-0.625** | **0.00142** | -0.360 | **0.01765** |
|  | Injured 72 hrs | Male |  | -0.031 | 0.89816 | **-0.597** | **0.00265** | -0.343 | **0.02437** |
|  | Injured 72 hrs | Female | Estradiol | **-0.555** | **0.01102** | -0.129 | 0.55654 | -0.251 | 0.10515 |
|  | Injured 72 hrs | Male |  | **-0.559** | **0.01045** | -0.106 | 0.63083 | -0.246 | 0.11229 |
| Angiogenesis involved in wound healing | Injured 24 hrs | Male | Androstenedione | -0.115 | 0.62807 | **-0.735** | **0.00006** | **-0.456** | **0.00213** |
|  | Injured 24 hrs | Male | Estradiol | **-0.618** | **0.00372** | -0.104 | 0.63763 | -0.257 | 0.09627 |
| Extracellular matrix organization | Injured 24 hrs | Female | Androstenedione | -0.094 | 0.69313 | **-0.713** | **0.00013** | **-0.439** | **0.00323** |
|  | Injured 72 hrs | Female |  | -0.083 | 0.72852 | **-0.692** | **0.00025** | **-0.425** | **0.00449** |
|  | Injured 24 hrs | Male |  | -0.099 | 0.67748 | **-0.710** | **0.00015** | **-0.440** | **0.00318** |
|  | Injured 72 hrs | Male |  | -0.079 | 0.74125 | **-0.692** | **0.00025** | **-0.423** | **0.00474** |
|  | Injured 24 hrs | Female | Estradiol | **-0.620** | **0.00354** | -0.139 | 0.52849 | -0.283 | 0.06623 |
|  | Injured 72 hrs | Female |  | **-0.618** | **0.00366** | -0.147 | 0.50315 | -0.285 | 0.06413 |
|  | Injured 24 hrs | Male |  | **-0.621** | **0.00351** | -0.141 | 0.52163 | -0.285 | 0.06369 |
|  | Injured 72 hrs | Male |  | **-0.620** | **0.00356** | -0.144 | 0.51239 | -0.285 | 0.06437 |
| Response to wounding | Injured 24 hrs | Female | Androstenedione | -0.101 | 0.67151 | **-0.731** | **0.00008** | **-0.440** | **0.00312** |
|  | Injured 72 hrs | Female |  | -0.083 | 0.72811 | **-0.719** | **0.00011** | **-0.432** | **0.00384** |
|  | Injured 24 hrs | Male |  | -0.101 | 0.67129 | **-0.726** | **0.00009** | **-0.439** | **0.00321** |
|  | Injured 72 hrs | Male |  | -0.074 | 0.75560 | **-0.712** | **0.00014** | **-0.427** | **0.00433** |
|  | Injured 72 hrs | Male | DHEA | **0.445** | **0.04920** | 0.040 | 0.85760 | 0.194 | 0.21280 |
|  | Injured 24 hrs | Female | Estradiol | **-0.621** | **0.00350** | -0.084 | 0.70447 | -0.268 | 0.08207 |
|  | Injured 72 hrs | Female |  | **-0.622** | **0.00343** | -0.102 | 0.64172 | -0.272 | 0.07770 |
|  | Injured 24 hrs | Male |  | **-0.616** | **0.00380** | -0.085 | 0.70051 | -0.266 | 0.08462 |
|  | Injured 72 hrs | Male |  | **-0.622** | **0.00339** | -0.093 | 0.67169 | -0.263 | 0.08831 |
| Tissue remodeling | Injured 24 hrs | Female | Androstenedione | -0.063 | 0.79072 | **-0.669** | **0.00048** | -0.383 | **0.01130** |
|  | Injured 72 hrs | Female |  | 0.151 | 0.52605 | **-0.607** | **0.00214** | -0.249 | 0.10678 |
|  | Injured 24 hrs | Male |  | -0.011 | 0.96230 | **-0.630** | **0.00126** | -0.337 | **0.02710** |
|  | Injured 72 hrs | Male |  | 0.151 | 0.52605 | **-0.607** | **0.00214** | -0.249 | 0.10678 |
|  | Injured 72 hrs | Male | DHEA | **0.460** | **0.04125** | 0.051 | 0.81390 | 0.219 | 0.15730 |
|  | Injured 24 hrs | Female | Estradiol | **-0.625** | **0.00319** | -0.072 | 0.74529 | -0.272 | 0.07791 |
|  | Injured 24 hrs | Male |  | **-0.603** | **0.00485** | -0.063 | 0.77461 | -0.258 | 0.09529 |

**Figure 2.**
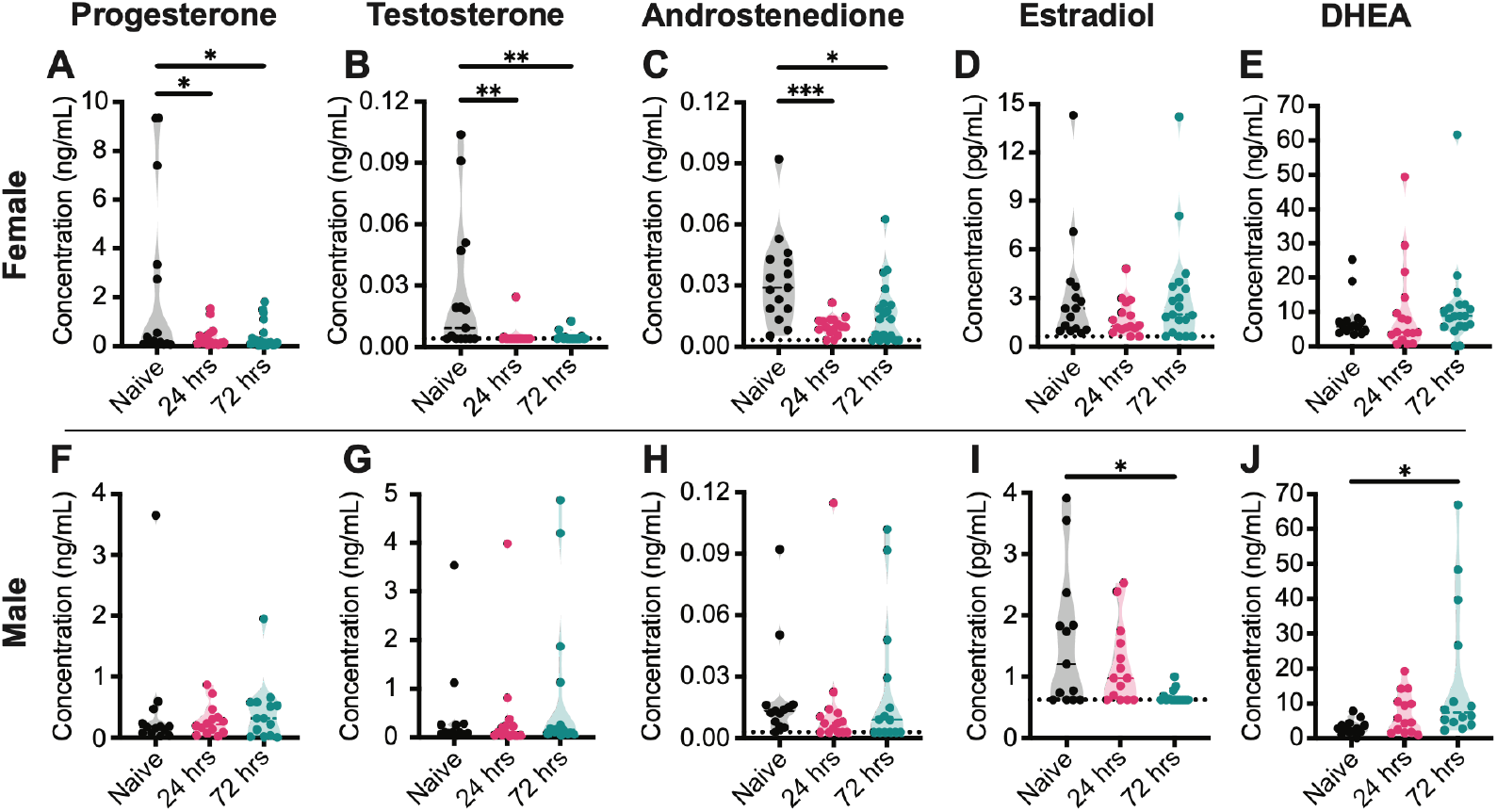
Circulating sex steroid hormone profile alterations in male and female mice after TBI. Quantification of progesterone, testosterone, androstenedione, estradiol and DHEA in female (**A-E**) and male (**F-J**) plasma samples. Dotted line shown on some graphs is the lower limit of detection for that analyte. One-way ANOVA, n=13-19 per cohort.(24 hrs = 24 hrs post-TBI, 72 hrs = 72 hrs post-TBI; *p<0.05, **p<0.01, ***p<0.001)

To determine sex differences in circulating steroid hormones across time-points, we utilized two-way ANOVAs to compare male and female levels of each hormone individually (Supplemental Figure 2; Supplemental Table 3,4). Circulating levels of progesterone (F=4.045, df=86, p=0.0475), testosterone (F=11.94, df=86, p=0.0009), and estradiol (F=11.1, df=86, p=0.0013) were significantly different between male and female groups across all timepoints (Sup. Fig. 2). Additionally, there was an injury main effect on DHEA (F=4.277, df=87, p=0.0170) and progesterone (F=4.05, df=87, p=0.0209).

**Table 3.** Results of Pearson correlation between eigengene vectors of genes that showed significant semantic similarity to tissue remodeling GO:CC terms and hormone levels for each cohort.

| GO:CC term | Timepoint | Sex | Hormone | Male Only |  | Female Only |  | All Samples |  |
| --- | --- | --- | --- | --- | --- | --- | --- | --- | --- |
|  |  |  |  | Male R | Male p-value | Female R | Female p-value | All R | All p-value |
| Adherens junction | Injured 24 hrs | Female | Androstenedione | -0.068 | 0.77465 | **-0.696** | **0.00023** | **-0.408** | **0.00658** |
|  | Injured 72 hrs | Female |  | -0.053 | 0.82350 | **-0.677** | **0.00039** | -0.396 | **0.00856** |
|  | Injured 24 hrs | Male |  | -0.073 | 0.76050 | **-0.693** | **0.00024** | **-0.409** | **0.00642** |
|  | Injured 72 hrs | Male |  | -0.053 | 0.82404 | **-0.676** | **0.00040** | -0.394 | **0.00903** |
|  | Injured 24 hrs | Female | Estradiol | **-0.610** | **0.00433** | -0.092 | 0.67617 | -0.272 | 0.07717 |
|  | Injured 72 hrs | Female |  | **-0.606** | **0.00463** | -0.107 | 0.62717 | -0.272 | 0.07723 |
|  | Injured 24 hrs | Male |  | **-0.607** | **0.00453** | -0.094 | 0.67026 | -0.272 | 0.07713 |
|  | Injured 72 hrs | Male |  | **-0.607** | **0.00454** | -0.101 | 0.64630 | -0.272 | 0.07789 |
| Extracellular matrix | Injured 72 hrs | Female | Androstenedione | 0.046 | 0.84599 | **-0.646** | **0.00087** | -0.351 | **0.02087** |
|  | Injured 72 hrs | Male |  | 0.043 | 0.85865 | **-0.647** | **0.00084** | -0.351 | **0.02106** |
|  | Injured 72 hrs | Female | Estradiol | **-0.621** | **0.00348** | -0.127 | 0.56460 | -0.262 | 0.08981 |
|  | Injured 72 hrs | Male |  | **-0.625** | **0.00318** | -0.118 | 0.59039 | -0.262 | 0.08928 |
| Tight junction | Injured 24 hrs | Female | Androstenedione | -0.069 | 0.77164 | **-0.687** | **0.00029** | **-0.412** | **0.00600** |
|  | Injured 72 hrs | Female |  | -0.063 | 0.79035 | **-0.667** | **0.00051** | **-0.404** | **0.00725** |
|  | Injured 72 hrs | Male |  | -0.065 | 0.78640 | **-0.664** | **0.00056** | -0.397 | **0.00843** |
|  | Injured 24 hrs | Female | DHEA | **0.455** | **0.04330** | 0.083 | 0.70766 | 0.224 | 0.14965 |
|  | Injured 72 hrs | Female |  | **0.451** | **0.04590** | 0.093 | 0.67238 | 0.224 | 0.14795 |
|  | Injured 72 hrs | Male |  | **0.448** | **0.04713** | 0.087 | 0.69306 | 0.224 | 0.14888 |
|  | Injured 24 hrs | Female | Estradiol | **-0.605** | **0.00470** | -0.130 | 0.55530 | -0.278 | 0.07063 |
|  | Injured 72 hrs | Female |  | **-0.589** | **0.00631** | -0.141 | 0.52141 | -0.271 | 0.07853 |
|  | Injured 72 hrs | Male |  | **-0.597** | **0.00544** | -0.125 | 0.57034 | -0.273 | 0.07656 |

### Sex differences in gene expression after TBI over time

We assessed temporal and sex-specific transcriptional changes to reveal molecular mechanisms and pathways disrupted after TBI (Supplemental Table 5). All comparisons were made between paired ipsi- and contra-lateral sides of each animal’s brain to account for batch effects (Figure 1C). Comparisons were made separately for each sex and timepoint (Figure 3A-F). As anticipated, there were no differentially expressed genes in naive animals for either sex (Figure 3A & D). An interesting sex-specific effect was observed within males whereby more significantly upregulated genes were present at 24 hrs, and fewer at 72 hrs post-TBI, whereas females had fewer upregulated genes at 24 hrs and more at 72 hrs post-TBI (Figures 3 B-C & E-F & G). Remarkably, a total of 485 upregulated genes were unique to the 72 hrs female mice, while only 72 upregulated genes were unique to the 24 hrs female mice (Figure 3G). The male mice had 227 and 190 genes uniquely upregulated genes across the 24 hrs and 72 hrs timepoints, respectively (Figure 3G). All four conditions had a similar amount of unique downregulated genes, with an average of 158 genes across both sexes at each timepoint (Figure 3H). These sex-specific differentially expressed genes indicate that there are differences in how each sex responds to TBI at the transcriptomic level, and exploring enriched biological processes or pathways may lead to further insights.

**Figure 3.**
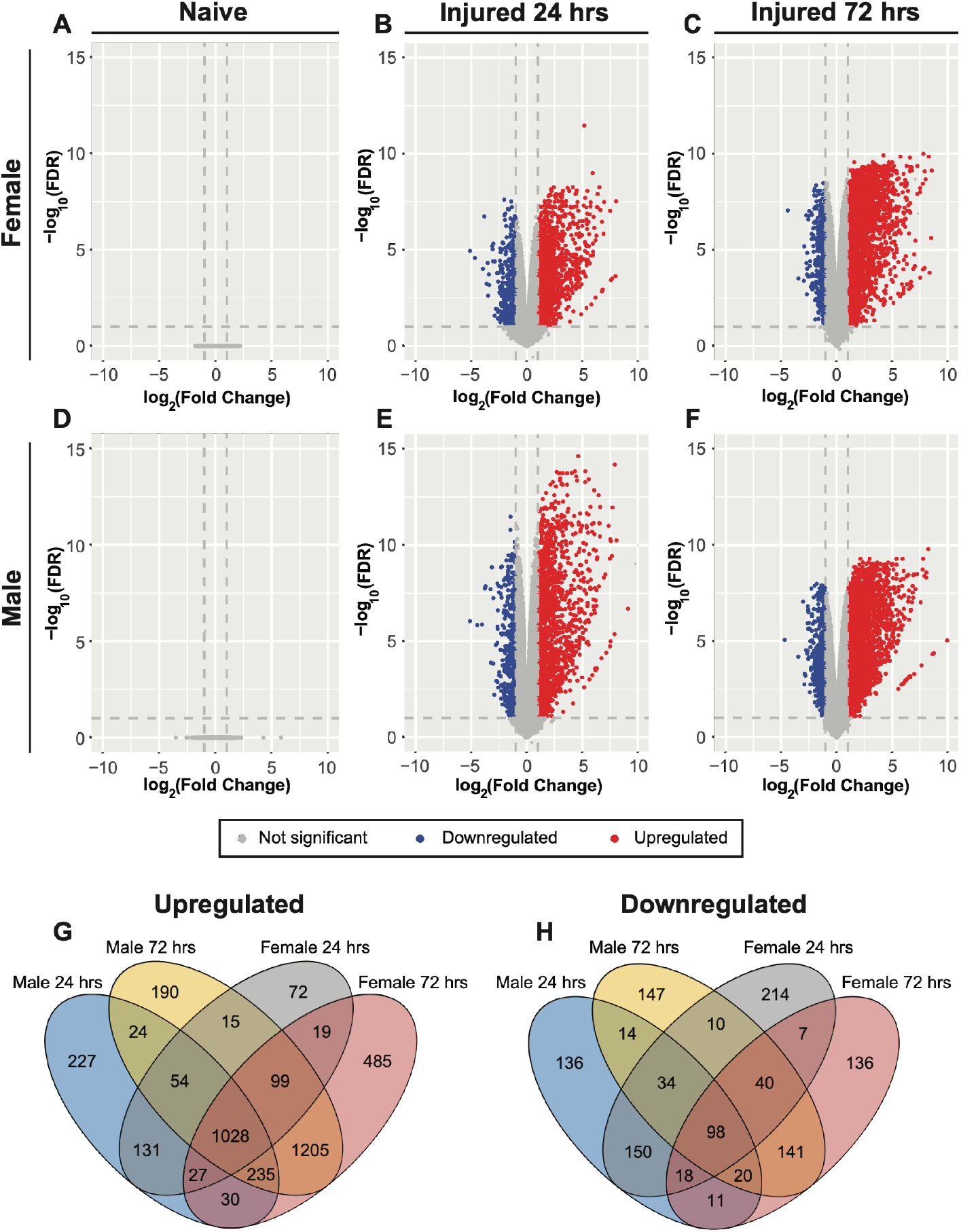
Sex differences in differential gene expression between ipsilateral and contralateral cortical tissue for male and female mice after TBI. Volcano plots showing differentially expressed genes (DEGs) naïve, injured 24 hr and injured 72 hr female (**A-C**) and male (**D-F**) cohorts (FDR: false discovery rate). Comparisons of male and female DEGs for each injured timepoint reveal unique gene sets (**G**,**H**).

### Characterizing the early innate immune response to cortical injury

The innate immune response becomes activated quickly after TBI, with substantial changes evident at acute timepoints (16). We characterized the activation of innate immune processes by performing functional enrichment analyses of the differentially expressed genes using Gene Ontology Biological Process (GO:BP) terms associated with innate immunity. We curated innate immune biological processes based on an overall term ‘innate immune response’, and more specific terms for ‘activation’ of innate immune cell types including macrophage, neutrophil, granulocyte, microglia, and three types of dendritic cells (follicular, plasmacytoid, and myeloid; Supplemental Table 6). Functional enrichment analysis was used to identify a list of GO:BP and GO:CC terms that are enriched for each sex and time point based differentially expressed gene set. Then, semantic similarity was used to identify associations between the functionally enriched GO:BP terms and the innate immunity curated terms (Supplemental Table 7).

Key players of the innate immune system, notably macrophage activation, neutrophil activation, and granulocyte activation, showed strong associations (Jiang-Conrath semantic similarity score > 0.8) with GO:BP terms enriched in cortical RNA-sequencing data at both 24 and 72 hrs post-TBI for each sex (Figure 4A). ‘Regulation of microglial cell activation’, an important component of the brain immune system, was also consistently associated at both timepoints for both sexes (Figure 4A). The sexes exhibited indistinguishable semantic similarity scores for these innate immune biological processes. We observed marked sex-specific differences in differential gene expression at each timepoint (Figure 4B-G). Within each timepoint, there are genes that have detectable gene expression for one sex and undetectable expression in the other (Figure 4 B-D). At 24 hrs post-TBI, 3 genes were expressed exclusively in females and 9 genes exclusively in males (Figure 4C). At 72 hrs post-TBI, 9 genes were expressed exclusively in females and 1 exclusively in males (Figure 4D). Many of these genes are involved in the production of proinflammatory cytokines and chemokines in response to injury, including *Scimp, Ppbp, Ccl8*, and *Treml1* (17–19). We also observed higher expression of *Saa1* in males at 24 hrs post-TBI, consistent with prior reports of sex-dependent Saa1 regulation after TBI (20). Furthermore, we observed groups of genes that exhibited higher expression in one sex relative to the other (fold change in expression > 2 between sexes) within each timepoint (Figure 4E-G). At 24 hrs post-TBI, males had a larger number of upregulated genes relative to females (Figure 4F). By 72 hrs, the differential expression becomes more similar across the sexes (Figure 4G). These differences in sex-specific gene expression and in the magnitude of expression describe the nuanced sex-dependent differences in immune responses to TBI.

**Figure 4.**
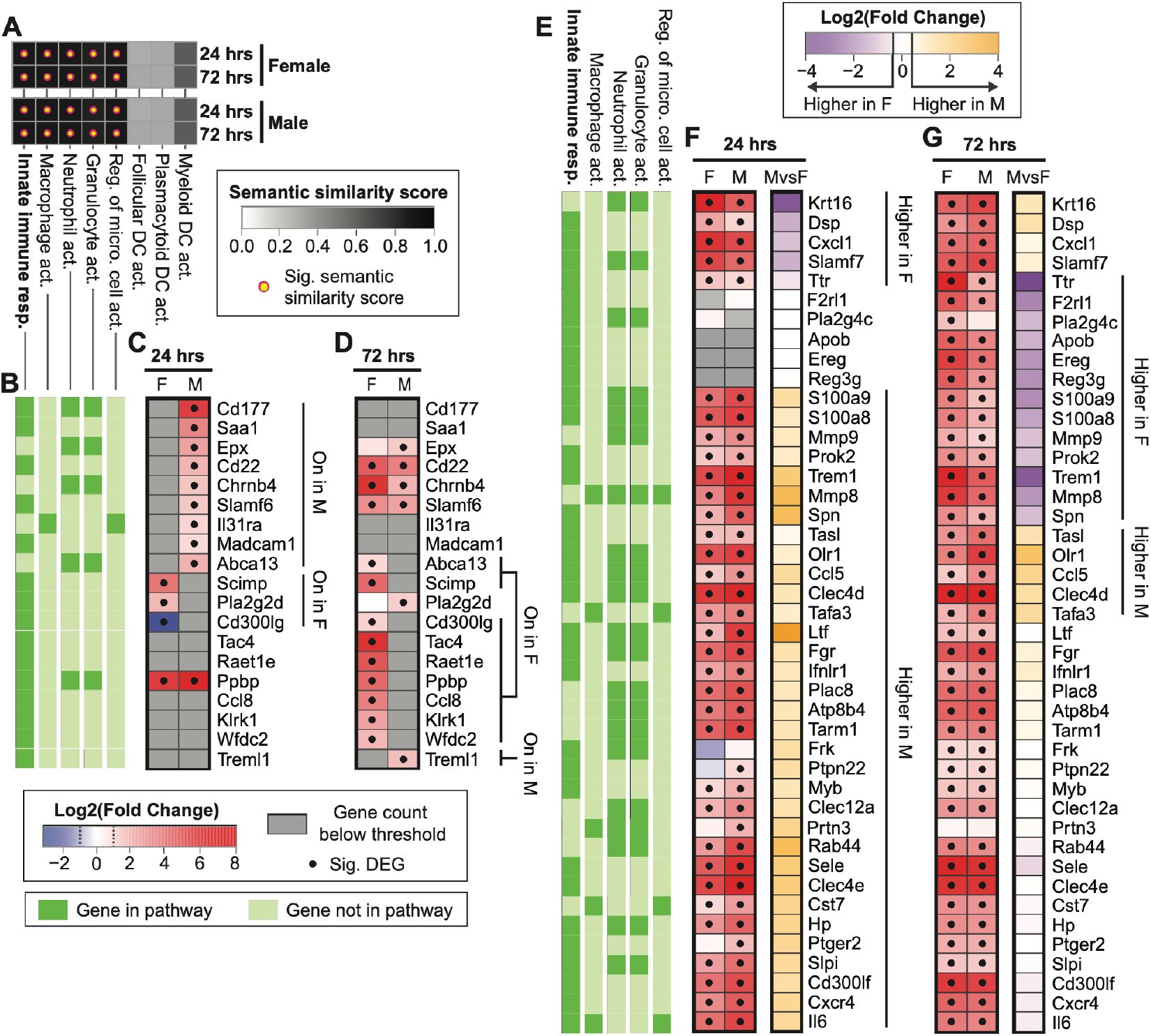
Sex differences in gene expression related to innate immune responses after TBI. Jiang-Conrath semantic similarity analysis was used to evaluate associations between curated innate immune GO:BP terms and functionally enriched GO:BP terms for each sex and time point based differentially expressed gene sets (**A**). Genes within pathways that showed significant associations were analyzed for sex differences in expression (**B-G**).

### Different sex hormones are correlated with similar immune responses in male and female mice

The expression of differentially expressed genes that contributed to associations with immune GO:BP terms were summarized into eigengene vectors and correlated with hormone levels. For injured female groups, androstenedione was negatively correlated with all significant immune processes from the semantic similarity analysis across both timepoints (Table 1), which is consistent with the known immunosuppressive action of androgens (21,22). For injured male groups, estradiol was negatively correlated with all significant immune processes in the semantic similarity analysis across both timepoints (Table 1), consistent with the known function of estradiol as a suppressor of neuroinflammation (23,24). Ultimately, although the same major innate immune pathways are engaged in both sexes following injury, we identify **sex-specific hormonal associations** with the biological processes underlying this response. These divergent hormone–immune relationships arise from distinct gene compositions within the enriched GO:BP terms, reflecting meaningful sex-dependent differences in the molecular architecture of the innate immune response.

### Characterization of tissue remodeling biological processes and cellular components after TBI

Remodeling of damaged tissue and surrounding extracellular matrix after insult is a complex process that involves many different cell types, including endothelial cells, astrocytes, oligodendrocytes, and microglia (25,26). This process begins promptly after injury as cells begin to release damage-associated molecular patterns (DAMPs) and can extend far past the initial insult. As such, we sought to determine if there was biological enrichment of tissue remodeling biological processes using semantic similarity analysis with GO:BP terms that were enriched with differentially expressed genes (Supplemental Table 8). Male and female cohorts exhibited significant semantic similarity scores (Jiang-Conrath semantic similarity score > 0.8) with the following GO:BP terms at both post-TBI timepoints: ‘extracellular matrix organization’, ‘response to wounding’, and ‘tissue remodeling’ (Figure 5A). At 72 hrs post-TBI, both sexes show significant semantic similarity scores with the ‘adherens junction organization’ GO:BP term. Additionally, males exhibited significant semantic similarity to ‘angiogenesis involved in wound healing’ at 24 hrs post-TBI. We observed sex differences in expression of genes within the semantically similar GO:BP terms (Figure 5B-G). We observe sex differences in the expression of multiple matrix metalloproteinase genes including *Mmp3, Mmp8, Mmp10, Mmp12, and Mmp13*, and *Mmp27*. Notably, these genes encode enzymes that promote neuroinflammation by breaking down extracellular matrix to promote infiltration of immune cells and by activating and functioning as neuroinflammatory signals (27). Furthermore, we observed sex differences in the expression of junctional protein genes *Cldn19* and *Tjp3*. Cadherins and claudins are critical proteins that are present at tight junctions and adherens junctions that comprise the BBB junction complex (28). These findings led us to explore the enrichment of GO Cellular Components (GO:CC) using semantic similarity analysis with GO:CC terms curated for cellular junctions (adherens junction, tight junction, extracellular matrix, desmosome, hemidesmosome, and gap junction; Figure 5H). We observed significant associations with ‘adherens junction’ and ‘tight junction’ GO:CC terms for females 24 hrs post-TBI, whereas males only exhibit significant associations with the ‘adherens junction’ GO:CC term at 24 hrs post-TBI. At 72 hrs post-TBI, male and female cohorts showed similar significant associations with the following GO:CC terms: ‘adherens junction’, ‘tight junction’, and ‘extracellular matrix’. Overall, these data indicate that while the same tissue-remodeling processes are engaged in both sexes following TBI, there are differences in the underlying transcriptional programs, providing a potential molecular basis for sex-dependent variation in injury outcomes.

**Figure 5.**
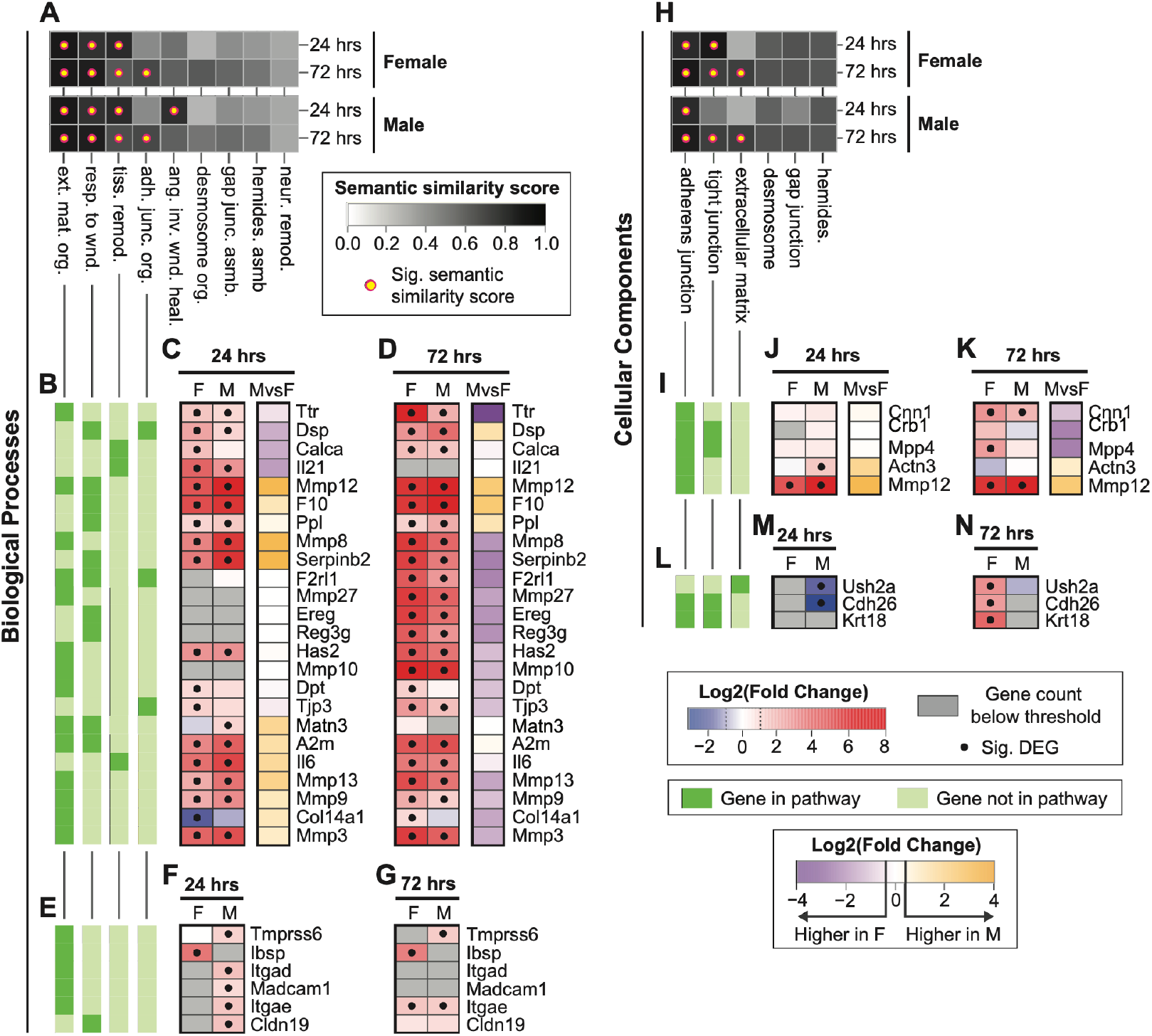
Sex differences in gene expression related to tissue remodeling after TBI. Jiang-Conrath semantic similarity analysis was used to evaluate associations between curated tissue remodeling GO:BP terms and functionally enriched GO:BP terms for each sex and time point based differentially expressed gene sets (**A**). Genes within pathways that showed significant associations were analyzed for sex differences in expression (**B-G**). Semantic similarity analysis was also used to evaluate associations between curated tissue remodeling GO:CC terms and functionally enriched GO:BP terms for each sex and time point based differentially expressed gene sets (**H)**. Genes within cellular components that showed significant associations were analyzed for sex differences in expression (**I-N**).

### Different sex hormones are correlated with similar tissue remodeling processes and cellular components in male and female mice

Next, eigengene vectors derived from genes within enriched tissue remodeling GO:BP and GO:CC terms were correlated with hormone levels. For injured female groups, we observe significant negative correlations between androstenedione level and all the significant tissue remodeling GO:BP and GO:CC terms in the semantic similarity analysis across both timepoints (Tables 2 & 3). Additionally, for injured male groups, we observed significant negative correlations between estradiol and all significant GO:BP and GO:CC terms across both timepoints (Table 2 & 3). We also observed significant positive correlations between DHEA and the following GO:BP and GO:CC terms: response to wounding (BP), tissue remodeling (BP), and tight junction (CC). Overall, these results describe sex-specific hormonal regulation of tissue-remodeling pathways and provide a potential a mechanism for a sex-dependent repair response.

### Vascular response: changes in junctional protein expression

Integration of transcriptomics and hormone levels identified genes that are likely to be vital for tissue remodeling. At 72 hrs post-TBI female differentially expressed genes were associated with the GO:CC term ‘Cell-Cell Junction’ which was significantly associated with the curated term ‘tight junction’ (Supplemental Table 9), and also associated with the hormone androstenedione (Table 3). Looking at genes from the enriched ‘Cell-Cell Junction’ term, we selected VECAD and CLDN5, as these proteins are critical for maintaining the integrity of the BBB (29,30). The endothelial cell marker CD31 enabled identification and delineation of blood vessel regions to assess the fluorescence intensity of VECAD and CLDN5 immunostaining specifically within blood vessels (Figure 6). Then, we evaluated the spatial distribution of these BBB junction complex proteins through immunostaining in the ipsilateral and contralateral hemispheres of male and female mice at 24 and 72 hrs post-TBI (Figure 5). Within the ipsilateral ROIs and CD31+ mask area, we identified significant effects with respect to injury of VECAD (F=10.72, df=21, p=0.0008), CLDN5 (F=9.705, df=21, p=0.0012) and CD31 (F=88.02, df=46, p<0.0001) mean fluorescence intensity (MFI; Figure 5U-W; Supplemental Tables 10, 11). Pairwise analysis revealed significantly increased VECAD MFI in the ipsilateral regions for the male injured 72 hrs group (t=3.085, df=19, p=0.0175) and the female injured 24 hrs (t=3.460, df=19, p=0.0079) and 72 hrs (t=2.883, df=19, p = 0.0286) groups compared to sex-matched naive controls. In contrast, pairwise analysis of CLDN5 MFI identified significant increased intensity in the male injured 24 hrs (t=2.766, df=19, p=0.0369) and 72 hrs (t=3.674, df=19, p=0.0048) groups in ipsilateral regions compared to sex-matched naive controls. For CD31 MFI, pairwise analysis indicated significant increases for both males and females at 24 hrs (male: t=5.827, df=44, p<0.0001, female: t=7.268, df=44, p<0.0001) and 72 hrs (male: t=9.07, df=44, p<0.0001, female: t=9.375, df=44, p<0.0001) post-TBI. Furthermore, there was a significant increase in CD31 intensity between 24 hrs and 72 hrs post-TBI for males (t=3.502, df=44, p=0.0032) and females (t=2.522, df=44, p=0.0461). In the contralateral region, the two-way ANOVA detected a significant effect with respect to injury for the VECAD intensity (F=3.894, df=21, p=0.0382) and CD31 (F=9.138, df=46, p=0.0005) (Figure 5X-Z; Supplemental tables 12, 13). Pairwise analysis revealed the only significant alteration in CD31 intensity between 24 hrs and 72 hrs post-TBI was in females (t=3.308, df=44, p=0.0056), with no significant effect in males. In parallel to the expression of these proteins, we also observed upregulation of their respective genes in ipsilateral cortices compared to contralateral cortices for all groups (Figure 5AA). Only *Cdh5* and *Pecam1* exhibited upregulation for both sexes at all timepoints, whereas *Cldn5* showed upregulation only at 72 hrs post-TBI for both sexes.

**Figure 6.**
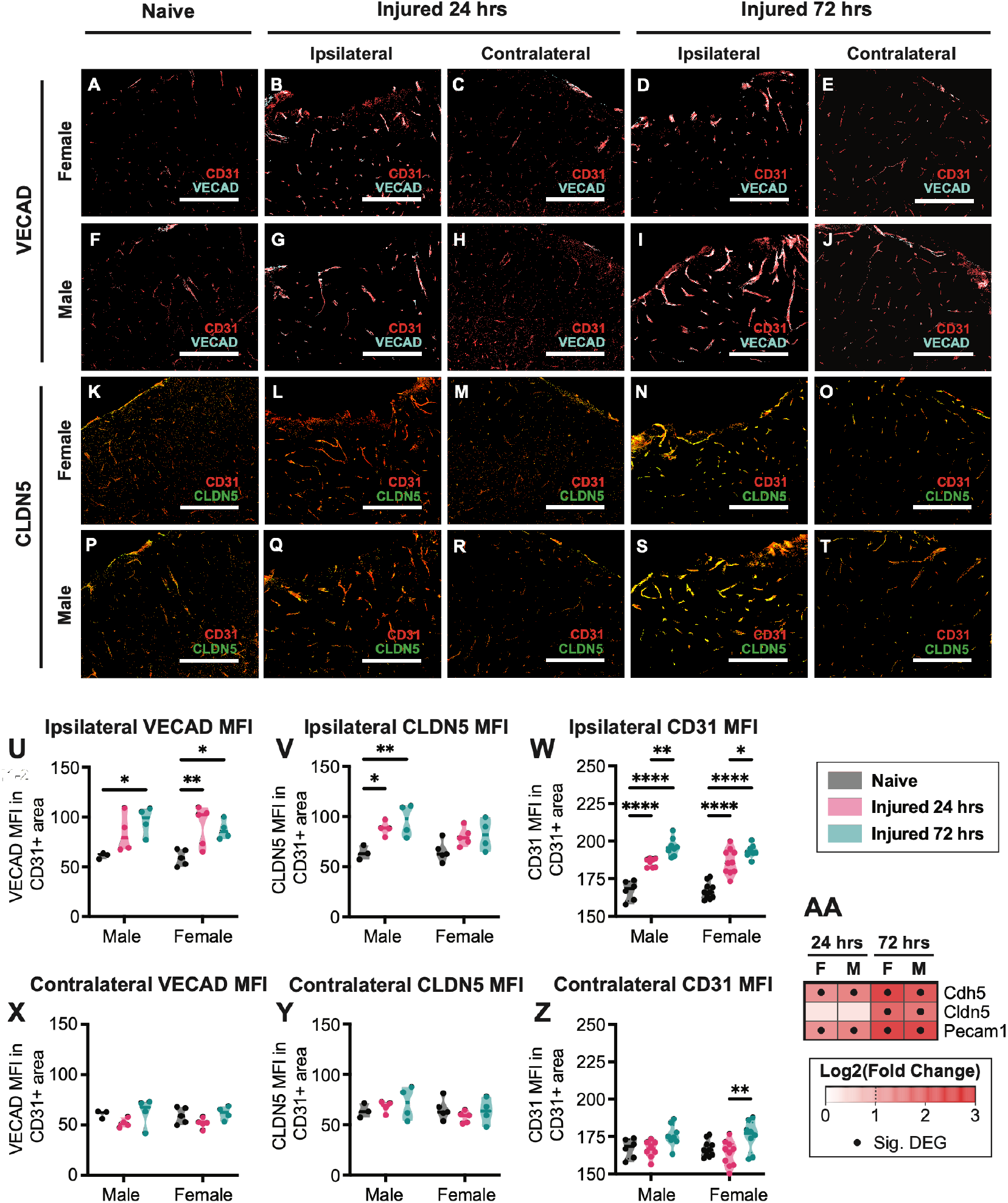
Evaluation of BBB junctional protein expression in response to CCI. Representative images of VECAD (**A-J**) and CLDN5 (**K-T**) expression in contralateral and ipsilateral regions of interest (ROIs) for male and female naïve, injured 24 hr, and injured 72 hr cohorts (scale bar = 200um). Quantification of VECAD, CLDN5, and CD31 staining in blood vessel area in ipsilateral ROIs for male and female cohorts (**U-W**). Quantification of VECAD, CLDN5, and CD31 staining in blood vessel area in contralateral ROIs for male and female cohorts (**U-W**). Visualization of expression of corresponding genes (Cdh5:VECAD, Cldn5:CLDN5, Pecam1: CD31) for injured male and female cohorts (**AA**). Two way ANOVA, mean ± SEM, n=3-5 per cohort.

## Discussion

Disruption in the HPA axis after TBI leads to hormonal dysregulation, yet the mechanism is not fully eluciated (31). The first step in unraveling underlying mechanisms is a robust characterization of the changes in circulating hormone profiles after TBI. This study directly addresses this knowledge gap by employing LC-MS/MS to characterize multiple circulating sex hormone levels after murine experimental brain injury in both male and female cohorts. Whereas previous studies have probed subsets of circulating hormone levels within a single sex cohort (10,32–34). Our data illustrate distinct alterations in circulating hormone profiles between male and female mice at two critical timepoints, both 24 and 72 hrs post-TBI. Injured females exhibited decreases in progesterone, testosterone, and androstenedione at both post-TBI timepoints, whereas injured males exhibited decreased estradiol and increased DHEA at 72 hrs post-TBI. Loss or reduction of endogenous progesterone, testosterone, and 17β− estradiol signaling has historically been linked to adverse consequences in the context of brain injury and this study stands as another testament (9,35). Moreover, we demonstrate significant correlations between the differential expression of genes enriched for secondary injury pathologies and the reduction of circulating steroid hormones.

Post-injury remodeling of damaged tissue is a dynamic process that involves interactions between local glial and endothelial cells, as well as infiltrating immune cells. These cells rapidly respond to DAMPs released by damaged tissue and initiate cascades of structural and molecular remodeling that can lead to neurodegeneration (16). Both sexes exhibited transcriptional changes for genes related to tissue remodeling and innate immune processes. However, despite these shared responses, further investigation revealed sex-based differential expression of genes that represented these processes. Differential expression of multiple matrix metalloproteinases and junctional proteins suggest sex differences in ECM degradation and BBB breakdown, which may impact the infiltration of immune cells. Further supporting this potential process, we observed sex differences in the expression of several genes involved in the production of pro-inflammatory signals. These findings support that although there are shared overarching responses to injury, the molecular restructuring of the ECM and BBB, as well as the resulting cellular immune infiltration is sex specific.

The primary limitation of this study is that we cannot decipher whether hormone levels, transcriptional changes, or the interactions between them, are driving our observed phenotypes. Previous studies have shown that manipulating steroid hormone levels impact the progression of secondary injury pathologies (36–38). Furthermore, it was not feasible to power our experiment to evaluate how the estrous cycle and its associated changes in sex hormone levels impact TBI pathology. Additional studies will be needed to determine how variations in hormone levels across the estrous cycle at the time of the TBI event could impact the trajectory of secondary injury pathologies for females.

Altogether, we demonstrated sex-specific, temporal alterations in circulating steroid hormone levels and cortical gene expression after TBI. This study provides a foundation for further investigation into the hormonal and transcriptional drivers of secondary injury. Future studies should investigate the underlying mechanisms linking hormonal to variations in molecular and cellular responses in the brain following injury. Importantly, further examination of how these sex-specific responses shape the trajectory of neuroinflammation, and other secondary injury mechanisms is crucial for the development of effective treatments for TBI.

## Materials and Methods

### Animals

All experiments involving animals were performed in accordance with established procedures approved by the Institutional Animal Care and Use Committee of Arizona State University (Protocol #23-1998R). Male and female 7-week-old C57BL/6J mice were purchased from Charles River Laboratories and housed under standard conditions. Upon arrival, mice were housed two per cage on ventilated racks on a 12-hr light/dark cycle. Mice had free access to a standard diet and water and were habituated for 7 days prior to experiments.

### Vaginal Cytology

After the habituation period, female mice were estrous cycle tracked via vaginal swab starting 10 days prior to injury (experimental groups) or sacrifice (naïve groups). Cycle tracking for injured female cohorts continued until sacrifice at designated timepoints. Male mice underwent similar handling to avoid confounding behavioral effects, similar to our prior control procedures (39). Cotton tipped swabs (Puritan PurSwab) were soaked in phosphate buffer solution (PBS) prior to conducting the vaginal smear. Smears were rolled onto glass slides and observed on a standard light microscope with 10x magnification. The estrous cycle phases was characterized as described in Goldman et al (40,41). Briefly, estrus was defined by predominantly anucleated cornified epithelial cells. Metestrus was defined by the presence of a combination of leukocytes, needle-like cells, and limited rounded epithelial cells. Diestrus was defined by the predominance of small round leukocytes with few other cell types. Proestrus was defined by primarily rounded nucleated epithelial cells, with some occasional leukocytes. Images were saved and evaluated for cycle phase classification at collection time by one researcher and validated by a blinded researcher at a later time.

### Cortical Controlled Impact

Adult 8–9-week-old mice (C57BL/6) underwent unilateral controlled cortical impacts (CCIs) over the right somatosensory cortex as previously described (12,42). Briefly, mice were anesthetized with 1.5-3% isoflurane and the surgical site was shaved. Mice were placed on a homeothermic warming system and secured in a mouse stereotactic frame. The surgical site was prepared with betadine and isopropyl prior to making a midline incision. A 3mm craniotomy was placed approximately 1mm lateral to the midline and 1mm posterior to bregma. The CCI was produced with a Leica Impact One CCI device using the following parameters: 2mm impactor tip, 6.0 m/s, 100ms duration, and 1mm depth. Any bleeding was controlled using cotton-tipped applicators and the incision was closed using 4-0 nylon sutures and subsequent topical application of a triple antibiotic ointment. Mice were given subcutaneous injections of buprenorphine (0.05 mg/kg) and sterile saline (0.5 mL) and moved to a clean warmed cage, where they were observed for 30-60 mins prior to returning to vivarium.

### Circulating steroid hormone analysis

At specified timepoints, mice received intraperitoneal injections of Euthasol (200 mg/kg). A thoracotomy was performed, and an EDTA-coated 23-gauge needle syringe was used to collect blood from the left ventricle; blood samples were stored on ice prior to processing. Blood samples were immediately centrifuged at 2000 RFC for 10 minutes at 4°C. After centrifugation, plasma was collected leaving the remaining cellular components undisturbed. Plasma samples were stored at −20°C prior to shipping to the Wisconsin National Primate Research Center for analysis. Steroid hormone analysis was conducted at the Assay Services Unit in the Wisconsin National Primate Research Center. The method was adapted from methods previously described (43,44). Briefly, internal standard was added to mouse plasma samples (100-200 μl) and then they were extracted twice using methyl tert butyl ether followed by dichloromethane. Samples were derivatized using dansyl chloride and then analyzed using SCIEX Triple Quadrupole Liquid Chromatographer with tandem Mass Spectrometer (Sciex 6500+ LC-MS/MS). Calibration curves were constructed for each analyte with at least 12 points. The linearity was r > 0.9990 and the curve fit was linear with 1/x weighting. None of the compounds of interest were detected in blank or double blank samples. Inter-assay coefficient of variation was determined by a pool of human plasma and a pool of mouse plasma and ranged from 5.6-9.7%.

### RNA-sequencing

Mice designated for RNA-sequencing (n=6-9 per sex per timepoint) were transcardially perfused with diethyl pryocarbonate-treated phosphate buffer solution (DEPC PBS) until liquid ran clear from the right atria. Brains were isolated and 4mm cortical biopsy punches centered over the injury site and the contralateral cortex were collected and placed in RNA protect solution (Qiagen RNAprotect Tissue Reagent). For naive animals, the cortical punch from the right hemisphere was labelled as ipsilateral, and the left cortical punch labelled as contralateral. Tissue samples were incubated in RNAprotect overnight at 4°C and then stored at −80°C for later processing. RNA isolation was performed on the cortical tissue using the Qiagen RNeasy Lipid Tissue Mini Kit. Samples were homogenized using Qiazol lysis reagent and 2mL dounce homogenizer. Chloroform was added and mixed, then the mixture was centrifuged to isolate the aqueous phase containing RNA. Purification was performed in accordance with kit procedures, except that centrifuge speeds were increased from 8,000 to 10,000 RPM during washes, and a final wash of the membrane was performed using 80% ethanol followed by a spin at max speed for 5 min using RNAse-free water to collect the purified RNA. Samples were shipped to Novogene Inc. for sequencing, quality control of raw reads, and genome alignment. The purified mRNA was isolated using poly-T magnetic beads, fragmented, and reverse transcribed into cDNA using random hexamer primers. Libraries were prepared through end repair, A-tailing, adapter ligation, size selection, PCR amplification, and purification. Quality and concentration of libraries were assessed by Qubit and real-time qPCR. Fragment size distribution was quantified using a bioanalyzer. Libraries were then pooled and sequenced on an Illumina platform, yielding ≥20 million reads per sample. Quality control of raw reads was performed to remove low-quality reads and reads containing adapter or poly-N sequences using an in-house Perl script. Clean RNA-sequencing data were aligned using Hisat2 v2.0.5 to the mouse genome mm10 (GCF_000001635.20). The resulting raw count files were used in downstream analyses.

### Differential gene expression analysis

Differential gene expression analysis was performed using the Bioconductor package edgeR between the ipsilateral and contralateral samples for each condition (45). The quasi-likelihood (QL) generalized linear model method was applied using a formulation that is a generalization of a paired t-test. This formulation detects genes that are differentially expressed between the ipsilateral and contralateral sides of the brain, adjusting for baseline differences between the mice (i.e., batch effects). Differentially expressed genes (DEGs) were considered significant if they passed a two-fold change (|log2(FC)| ≥ 1) threshold in either regulation direction and a Benjamini-Hochberg corrected hypothesis false discovery rate cutoff of less than or equal to 0.1.

### Determining biological enrichment of tissue remodeling and the immune system during traumatic brain injury using RNA expression data

We used the search terms ‘immune system’ and ‘tissue remodeling’ to manually curate Gene Ontology Biological Process (GO:BP) and Gene Ontology Cellular Components (GO:CC) terms that describe biology impacted by traumatic brain injury (Supplemental Table 6).

Independently, functional enrichment analyses were performed on the differentially expressed gene sets to identify enriched GO:BP (GO_Biological_Process_2021) and GO:CC (GO_Cellular_Component_2025) terms using enrichr v0.14.0 (GSEApy). GO terms were considered significant if they had a Benjamini-Hochberg adjusted p-value cutoff of 0.05 and a gene overlap of at least 10%. The Jiang-Conrath semantic similarity score was used to link enriched GO:BP and GO:CC terms with the curated sets of terms we defined using the R package GOSemSim v3.16 (46). A Jiang-Conrath semantic similarity score of at least 0.8 was considered significant for the biological enrichment of terms describing the immune response and tissue remodeling (47). The genes from semantically similar terms were identified within the same timepoint as genes that were either on in only one condition (i.e., expressed in one sex and below the read count cutoff for the other), or significantly upregulated in one sex over the other (i.e., ≥ 2 fold-change difference).

### Correlating cortical gene expression with steroid hormones using differentially expressed eigengenes

Gene counts were normalized using variance stabilized transformation (vst) in DESeq2 (48). Batch effects were removed by normalizing gene expression of ipsilateral sample to the contralateral sample from the same mouse brain. Eigengenes were used to summarize the common variation across differentially expressed gene sets present in enriched GO:BP and GO:CC terms that were semantically similar to the curated GO terms defining immune system and tissue remodeling biology. An eigengene is the first principal component of the DEGs for a condition corrected for sign (49). Each eigengene was correlated with each steroid hormone across male and female samples using a Pearson correlation the pearsonr function in the scipy.stats package (50). Correlations were considered significant if the p-value was less than or equal to 0.05 and the absolute value of the Pearsons correlation coefficient was greater than 0.4.

### Immunohistochemistry

Mice designated for immunohistochemistry (n=3-5 per sex per timepoint) were transcardially perfused with ice cold PBS followed by a 4% paraformaldehyde solution. Brains were collected and placed in 4% paraformaldehyde solution overnight at 4°C prior to immersion in sequential incubations with 15% and 30% sucrose solutions. Super cooled ethanol was used to freeze fixed brains in Optimal Cutting Temperature medium (OCT). Frozen brain samples were stored at –80°C until coronally cyrosectioned at a thickness of 30 μm (Leica CM1950 cryostat). Brain tissue sections were permeabilized with 0.3% Tween-20 PBS, then blocked with 5% horse serum in 0.01% Tween-20 PBS. Sections were stained for their respective primary and secondary antibodies according to Supplemental Table 14. Upon completion of antibody staining, DAPI (NucBlue Fixed Cell ReadyProbes Reagent, ThermoFisherScientific, R37606) was added to stain cell nuclei. Sections were mounted onto charged glass slides (Globe Scientific, 1358W), coverslipped in Fluoro-Gel (Electron Microscopy Sciences, 17985-41), and stored at 4°C prior to imaging.

### Confocal Imaging and quantification

Tissue sections were imaged at 20X magnification using a Zeiss LSM 800 laser scanning confocal microscope (LSCM). All images were collected with the same acquisition settings to allow for direct comparisons between tissue sections. Five sequential sections per animal for each of the CLDN5 and VECAD stains were analyzed starting at approximately −0.5 mm bregma within the somatosensory cortex. The Indica Labs HALO Image Analysis software was used to quantify vascular endothelial cell expression of CLDN5 and VECAD in the injury penumbra. The ipsilateral region of interest (ROI) was defined as a 500 μm by 200 μm area starting at the most lateral edge of the injury moving medially along the injury penumbra. The pen tool was used to annotate 500 μm of the injury edge, which was copied 200 μm below to define the area. For contralateral and naive regions, a 500 μm by 200 μm rectangular area within layers I-IV of the primary somatosensory cortex was analyzed. The area quantification analysis was defined by the CD31+ vasculature (pixel intensity > 145); this threshold removed background noise and was held constant for all images. For both the CLDN5 and VECAD channels, the mean fluorescence intensity (MFI) was quantified within the CD31+ area. Data are presented as MFI means from 3-5 biological replicates; each biological replicate consisted of MFI means over at least 5 tissue sections. Representative images of ROIs were cropped in Adobe Photoshop.

### Statistical analysis

Statistical analyses of hormone and immunohistochemistry data were conducted in GraphPad Prism 10 (GraphPad Software, Inc.) using ordinary one-way and two-way ANOVAs. If the effect of timepoint or sex was significant (p<0.05), then post hoc analysis was performed using Bonferroni’s multiple comparison test.

## Acknowledgments

We thank Dr. Amita Kapoor and Cody Corbett in the Assay Services Facility at the Wisconsin National Primate Research Center for the LC-MS/MS services and collaboration; ASU Regenerative Medicine Core Facility for providing and maintaining equiptment and resources used to process tissues with the IHC analysis; Scott Stevens and Alondra Davila for assisting with experiments in the laboratory; All our laboratory members for their insights, support and collaboration.

## Author contributions

Conceptualization: SES, HBN, RWS, CP

Methodology: SES, HBN, RWS, JN, CP, AS, SW, VP

Investigation: SES, CP, AS, SW, SC

Visualization: SES, CP, AS, SW

Supervision: SES, HBN, RWS, CP, JN

Writing—original draft: SES, CP, AS, SW

Writing—review & editing: SES, HBN, RWS, JN, CP, AS, SW

## Supplementary Materials

Supplemental figures 1-2 and supplemental tables 1-14 in separate files.

