## Supplementary figures and images for "Differential sex-dependent responses of circulating steroid hormones and cortical gene expression in a preclinical traumatic brain injury model"

### Supplemental Figure 1

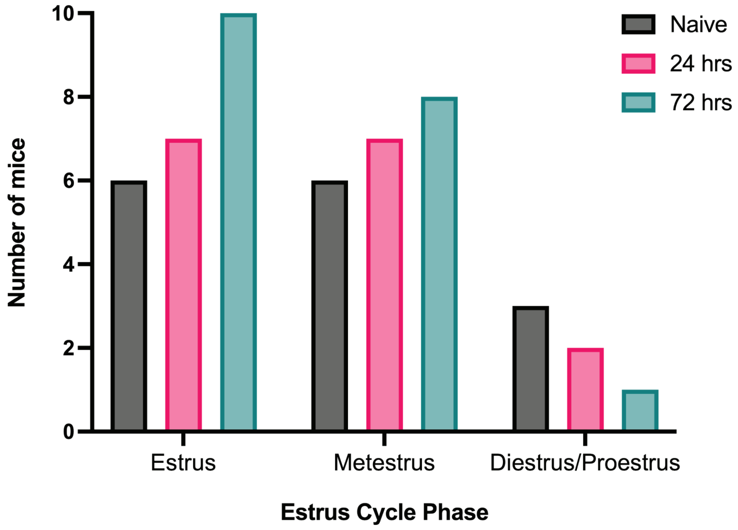

### Supplemental Figure 2

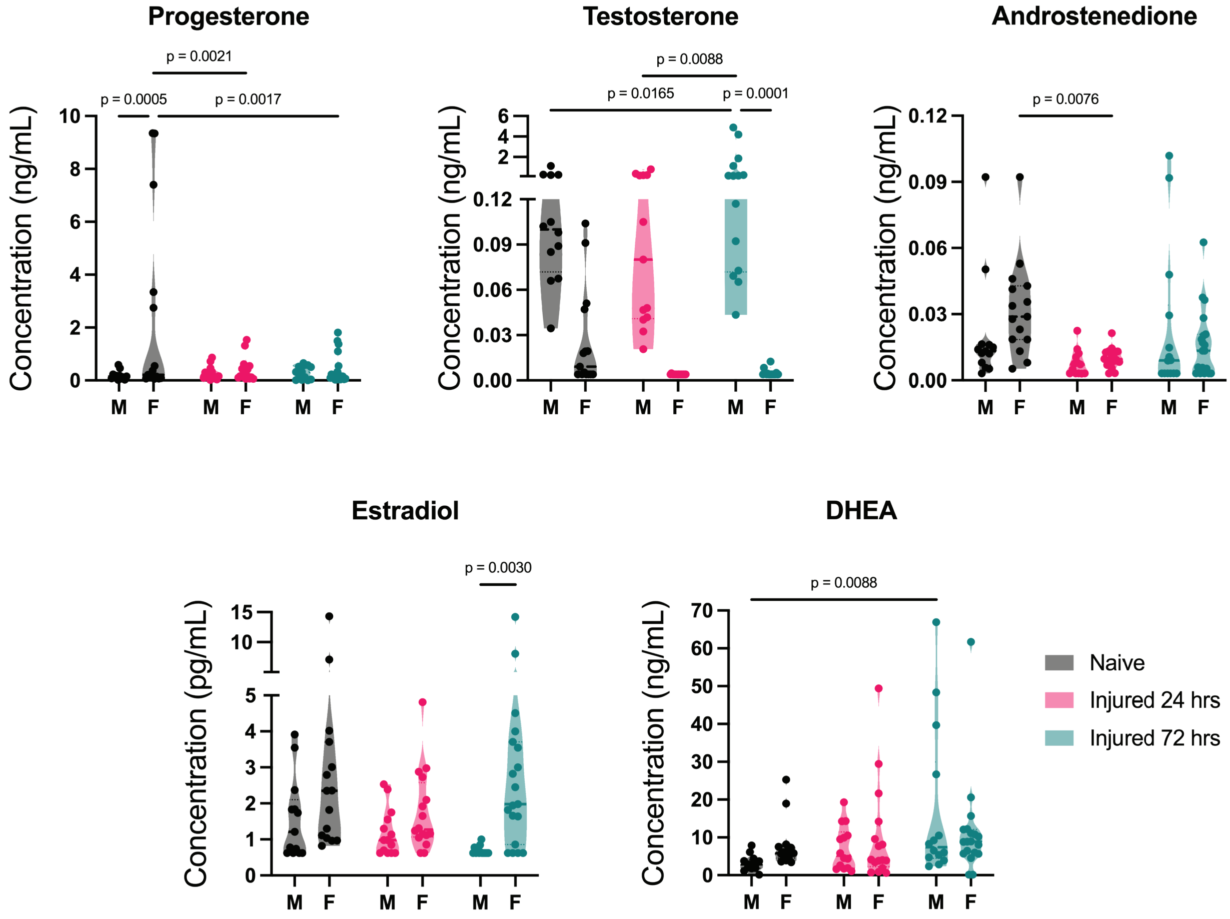
